# Total virome characterizations of game animals in China reveals a spectrum of emerging viral pathogens

**DOI:** 10.1101/2021.11.10.467646

**Authors:** Wan-Ting He, Xin Hou, Jin Zhao, Jiumeng Sun, Haijian He, Wei Si, Jing Wang, Zhiwen Jiang, Ziqing Yan, Gang Xing, Meng Lu, Marc A. Suchard, Xiang Ji, Wenjie Gong, Biao He, Jun Li, Philippe Lemey, Deyin Guo, Changchun Tu, Edward C. Holmes, Mang Shi, Shuo Su

**Affiliations:** Jiangsu Engineering Laboratory of Animal Immunology, Institute of Immunology, College of Veterinary Medicine, Nanjing Agricultural University, Nanjing, China; The Centre for Infection and Immunity Studies, School of Medicine, Sun Yat-sen University, Guangzhou, China; Changchun Veterinary Research Institute, Chinese Academy of Agricultural Sciences, Changchun, China; MOA Key Laboratory of Animal Virology, Zhejiang University, Hangzhou, China; Department of Biostatistics, Fielding School of Public Health, and Departments of Biomathematics and Human Genetics, David Geffen School of Medicine, University of California Los Angeles, Los Angeles, CA, the United States; Department of Mathematics, School of Science & Engineering, Tulane University, New Orleans, LA, USA; Agricultural College, Jinhua Polytechnic, Jinhua, China; Department of Infectious Diseases and Public Health, Jockey Club College of Veterinary Medicine and Life Sciences, City University of Hong Kong, Hong Kong, China; Department of Microbiology, Immunology and Transplantation, Rega Institute, Laboratory for Clinical and Epidemiological Virology, KU Leuven, Leuven, Belgium; Sydney Institute for Infectious Diseases, School of Life and Environmental Sciences and School of Medical Sciences, The University of Sydney, Sydney, NSW 2006, Australia

**Author notes:** Corresponding authors: Shuo Su; Mang Shi; and Edward C. Holmes. These authors contributed equally (Wan-Ting He, Xin Hou, Jin Zhao).

## Abstract

Game animals are wildlife species often traded and consumed as exotic food, and are potential reservoirs for SARS-CoV and SARS-CoV-2. We performed a meta-transcriptomic analysis of 1725 game animals, representing 16 species and five mammalian orders, sampled across China. From this we identified 71 mammalian viruses, with 45 described for the first time. Eighteen viruses were considered as potentially high risk to humans and domestic animals. Civets (*Paguma larvata*) carried the highest number of potentially high risk viruses. We identified the transmission of *Bat coronavirus* HKU8 from a bat to a civet, as well as cross-species jumps of coronaviruses from bats to hedgehogs and from birds to porcupines. We similarly identified avian *Influenza A virus* H9N2 in civets and Asian badgers, with the latter displaying respiratory symptoms, as well as cases of likely human-to-wildlife virus transmission. These data highlight the importance of game animals as potential drivers of disease emergence.

**Highlights:**

- 1725 game animals from five mammalian orders were surveyed for viruses
- 71 mammalian viruses were discovered, 18 with a potential risk to humans
- Civets harbored the highest number of potential ‘high risk’ viruses
- A species jump of an alphacoronavirus from bats to a civet was identified
- H9N2 influenza virus was detected in a civet and an Asian badger
- Humans viruses were also identified in game animals

## Introduction

Mammalian game animals are wildlife or semi-wild animals that are commonly traded and consumed as exotic food in China and other Asian countries^1–3^.They include rodents (such as porcupines, bamboo rats and marmots), carnivores (such as civets, badgers and foxes), pangolins, hedgehogs and rabbits. These animals are normally either caught and raised locally or imported illegally from neighboring countries before being transferred to live animal (or “wet”) markets for trading^1,3–7^, and in recent decades there has been a major expansion in commercial wildlife farming operations and their species diversity. The Huanan Seafood Wholesale Market in Wuhan, to which many of the early COVID-19 cases were linked^8^, is a notable example of a live animal market. Poor hygiene conditions and close contact between animals and humans, as well as a wide mix of species within live animal markets and the restaurants they serve, make them ideal breeding ground for emerging infectious diseases.

Unsurprisingly, consuming, capturing, processing and/or trading game animals has been linked to several important infectious disease outbreaks with grave public health consequences. Early cases of both SARS-CoV and SARS-CoV-2 were identified in animal handlers at animal markets in Guangdong^9^ and Hubei provinces^8^, respectively, and close relatives of SARS-CoV and SARS-CoV-2 have been identified in civets^10^, raccoon dogs^10^, and pangolins^11–13^. These are the most popular exotic game animals and subject to frequent trading and human consumption^14^, although their role as direct hosts for the transmission of these viruses to humans remains to be confirmed. In an analogous manner, *Human immunodeficiency virus* type I (HIV), the causative agent of AIDS, likely originated from the hunting or handing of carcasses of common chimpanzees in central-west Africa^15–17^.

As game animals are frequently associated with important human diseases, it is of obvious importance to identify existing or potential pathogens within these species so they can be used to trace the origins of specific epidemics and provide a risk assessment of the most likely sources of future outbreaks. Since the first SARS-CoV outbreak in 2002/2003, virus discovery studies have been performed in a variety of game animal species including civets, pangolins, marmots, and badgers. Initially, these studies utilized virus isolation, consensus PCR and Sanger sequencing approaches^10,18–23^, while in recent years more attention has been directed toward metagenomic next-generation sequencing (mNGS)^24–27^. In addition to SARS-like viruses, these studies have discovered several infectious agents that are of direct importance for human infection, such as *Rotavirus A* from civets and raccoon dogs^28,29^ and *Hepatitis E virus* in wild boar^30^. Both of these viruses are known to infect humans along with a wide range of mammalian hosts^31–33^. Despite this, there have been few systematic investigations of the virome in game animals, especially in China where their consumption is commonplace. For example, one study identified a virus commonly associated with pneumonia in rodents - *Sendai virus-* in pangolins, suggesting the epizootic potential of viruses in game animals^24^.

Since the emergence of SARS-CoV in 2002 China has been the focus of intense viral surveillance in wildlife animals, and a number of SARS-CoV-2 related viruses have been discovered in bats^34,35^. However, with the exception of pangolins, there has been little investigation of game animals, even though they have close contact with humans and domestic animals and hence provide a link to other wildlife species. To help fill this gap we performed a systematic meta-transcriptomic (i.e. total RNA sequencing) virome investigation of 16 species of game animals representing five mammalian orders collected across China. Many of the species were investigated for the first time using a metagenomic framework. Our purpose was to reveal the diversity and abundance of vertebrate-associated viruses in these game animals and assess which species have the greatest potential for carrying viruses that could eventually emerge in human populations.

## Results

### Game animals studied

Between 2017 and 2021 we performed a large-scale survey of viral pathogens in game animals commonly consumed as exotic food throughout China (Figure 1A). Of these animal samples, 98.55% (1700/1725) were collected after February 2020 and hence concurrent with the COVID-19 pandemic. Only a subset of the pangolin samples (n = 11) were collected between 2017 to 2019.

**Figure 1.**
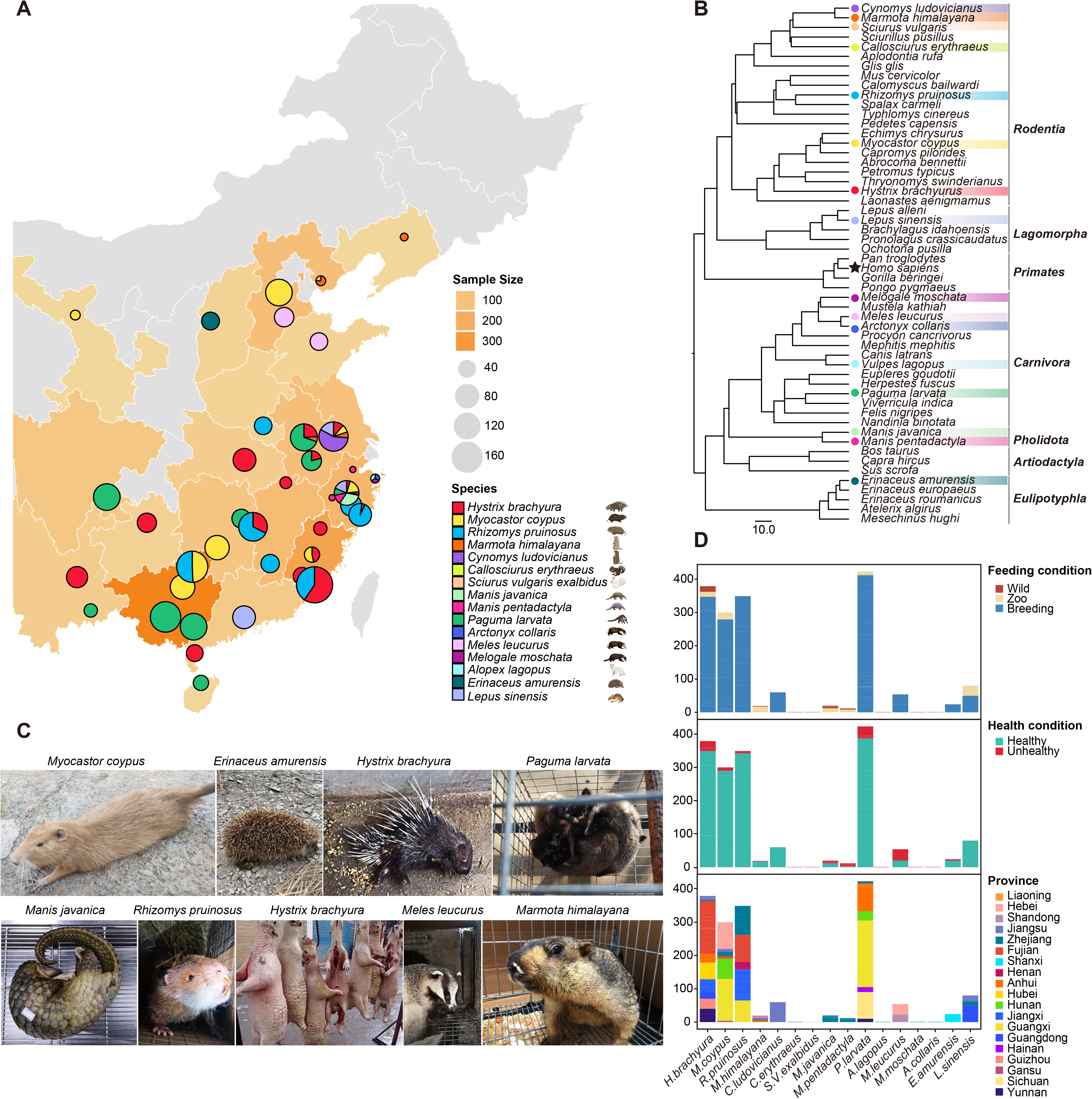
Game animals analyzed in this study. (A) Circles on the map show the geographical locations in China where the game animals were surveyed (n=2595 samples from 1725 individuals) during the period June 2017 to June 2021. Circle colors indicate the animal species according to the legend, and the Chinese provinces sampled are shown in shades of yellow. (B) Phylogenetic relationships of the game animal hosts surveyed here and related representative mammalian species. Circles denote the species included in this study, and stars highlight the phylogenetic position of humans. (C) Pictures of representative animals surveyed here. (D) Distribution of game animal samples by feeding condition (top panel), health condition (middle panel) and sampling province (bottom panel).

The total data comprised 1725 animals from 16 species representing five mammalian orders: *Rodentia*, *Pholidota*, *Carnivora*, *Eulipotyphla* and *Lagomorpha* (Figure 1B), including civets (*P. larvata*) that have been implicated in the emergence of SARS-CoV^36^. While most of these animals were maintained in artificial breeding sites that supply animal markets and zoos (Figure 1D, upper panel), some were obtained from their natural habitats. In addition, some animals had obvious signs of infectious disease and some even died during collection (Figure 1D, middle panel), presenting with such symptoms as paralysis in porcupines, anorexia and convulsion in pangolins, and influenza-like symptoms (nasal excretions) in Himalayan marmots (Figure 1C). Other animals exhibited no overt signs of disease. Respiratory and fecal samples were collected from these animals across 19 provinces in China (Figure 1D, bottom panel), which were subsequently organized into 181 pools according to species, location, health and living condition for meta-transcriptomic sequencing. This process yielded 172.36 billion nucleotide bases of sequence reads for virus discovery and characterization.

### Virome characterization

Despite the very large number of viruses discovered, we focused on those associated with vertebrates, comprising: (i) vertebrate-specific viruses that exhibited relatively close phylogenetic relationships to virus families or genera already known to infect vertebrates, and (ii) vector-borne viruses previously associated with both vertebrates and arthropods. Other viruses likely associated with animal diet, co-infecting parasites, or endosymbionts, were not considered further.

A total of 71 vertebrate-associated viral species from 13 viral families were identified based on meta-transcriptomic sequencing and confirmed by RT-PCR and Sanger sequencing (Figure 2A). This comprised six species of DNA virus belonging to the genera *Dependoparvovirus*, *Bocavirus*, *Protoparvovirus* and *Chapparvovirus* of family *Parvoviridae* and 65 RNA virus species belonging to the genera/families *Picornaviridae*, *Astroviridae*, *Paramyxoviridae*, *Orthomyxoviridae*, *Orthopneumoviruis*, *Flaviviridae*, *Reoviridae*, *Coronaviridae*, *Caliciviridae*, *Tobaniviridae*, *Hepeviridae* and *Birnaviridae* (Figure 2B). Among these, viruses of the *Picornaviridae*, *Astroviridae* and *Flaviviridae* were the most commonly detected and showed relative high abundance and prevalence in many animal species, whereas other viral families were more sporadically detected (Figure 2C). For example, coronaviruses were identified in bamboo rats, civets and hedgehogs, influenza viruses were identified in civets and Asian badgers, caliciviruses were identified in bamboo rats, rabbits, civets and Asian badgers, and the genera *Orthorubulavirus* (*Paramyxoviridae*) and *Pestivirus* (*Flaviviridae*) were mainly identified in pangolins (Figure 2C). Importantly, no viruses closely related to either SARS-CoV or SARS-CoV-2 (or other sarbecoviruses) were detected in any of animals examined.

**Figure 2.**
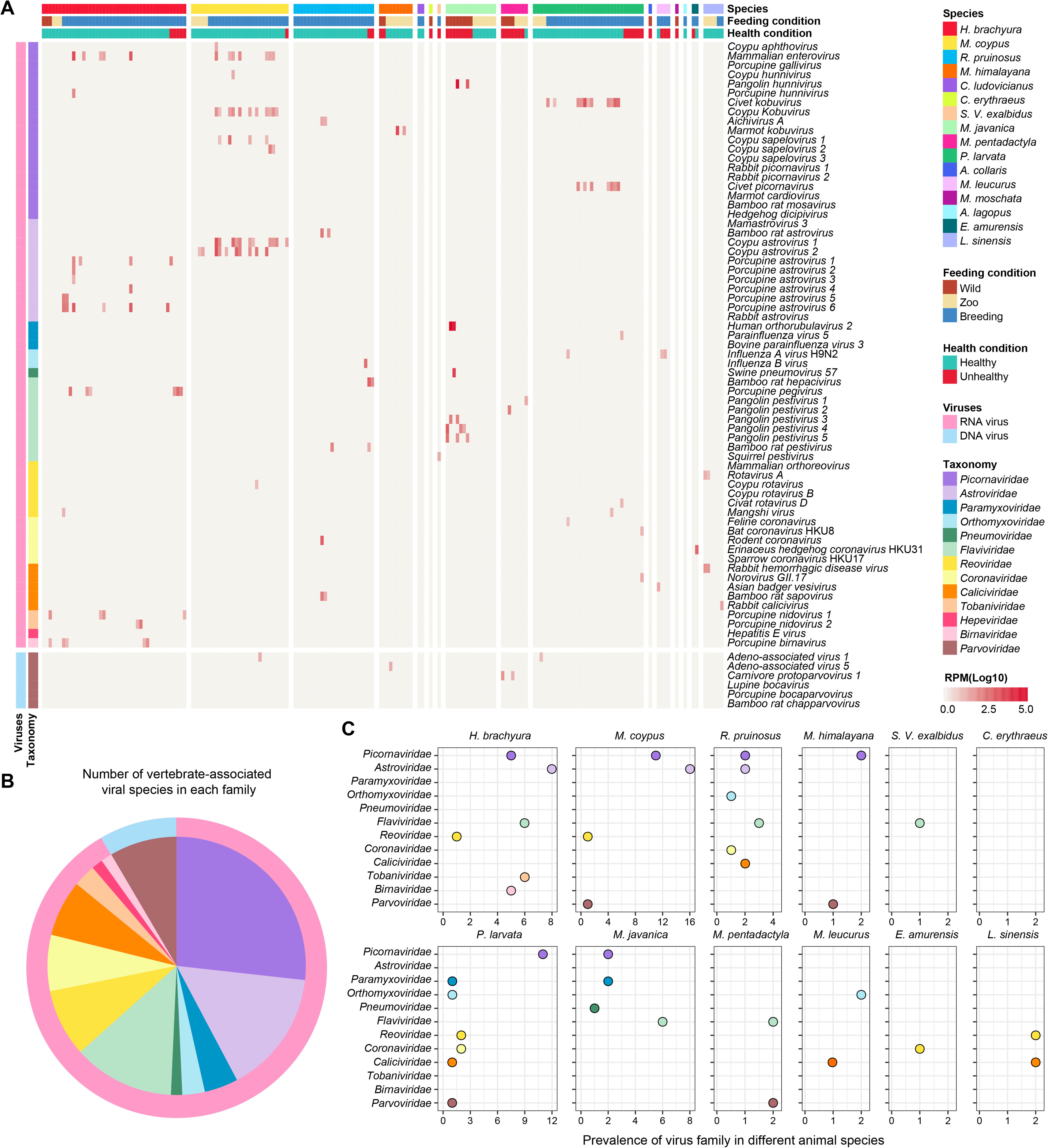
Overview of game animal viromes. (A) The distribution and abundance of vertebrate-associated viruses in game animal species. The relative abundance of viruses in each library was calculated and normalized by the number of mapped reads per million total reads (RPM). To remove potential contamination, only viruses with an abundance RPM > 10 are shown. Animal feeding and health condition were recorded when sampling and reflected in the corresponding colors. Viral species from 13 families are shown, with each family indicated by the colors on the heatmap. (B) Number of vertebrate-associated viral species in each family. (C) Prevalence (i.e., number of positive libraries) of each viral family in the animal species surveyed. Species with no vertebrate-associated viruses are not shown.

### Evolutionary history of vertebrate-associated viruses

Phylogenetic analyses of the viruses identified here revealed that many had close evolutionary relationship (>80% nucleotide sequence identity) to virus species known to cause disease in other wildlife species, domestic animals or even humans, thereby greatly expanding their host range (Figure 3 and 4, Figure S1). In particular, viruses associated with human infection, namely *Influenza A virus* (subtype H9N2), *Influenza B virus*, *Norovirus GII.17*, *Hepatitis E virus* (HEV), *Human parainfluenza virus 2* (HPIV2), *Rotavirus* A and *Mammalian orthoreovirus*, were identified in game animals for the first time. Notably, we report the first identification of *Influenza A virus* H9N2 and HEV in badgers and civets, respectively, while HPIV2, *Norovirus GII.17* and *Influenza B virus*, previously thought to be human-specific, were detected in pangolins, civets and bamboo rats at moderate or high abundance (40.01-293198.54 reads per million total reads (RPM)).

**Figure 3.**
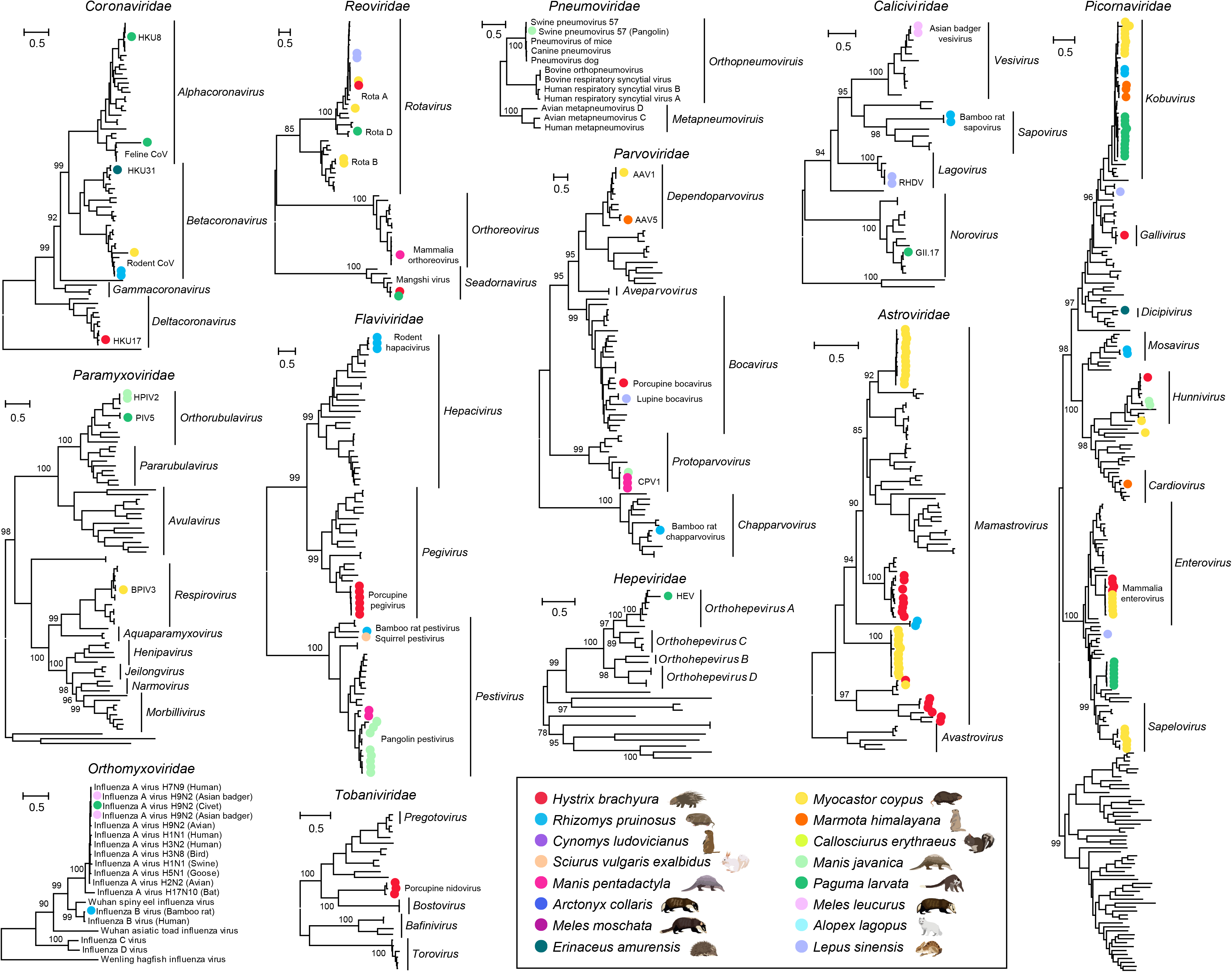
Inter-specific phylogenetic relationships of 12 major vertebrate-associated virus families. Each phylogenetic tree was estimated using a maximum likelihood method based on conserved viral proteins (RNA viruses = RdRp domain; DNA viruses = major capsid protein). All trees were midpoint-rooted for clarity only, and the scale bar indicates 0.5 amino acid substitutions per site. Bootstrap values are shown for major nodes. Within each phylogeny, the viruses newly identified here are marked by with solid circles and colored by animal host as shown in the key.

**Figure 4.**
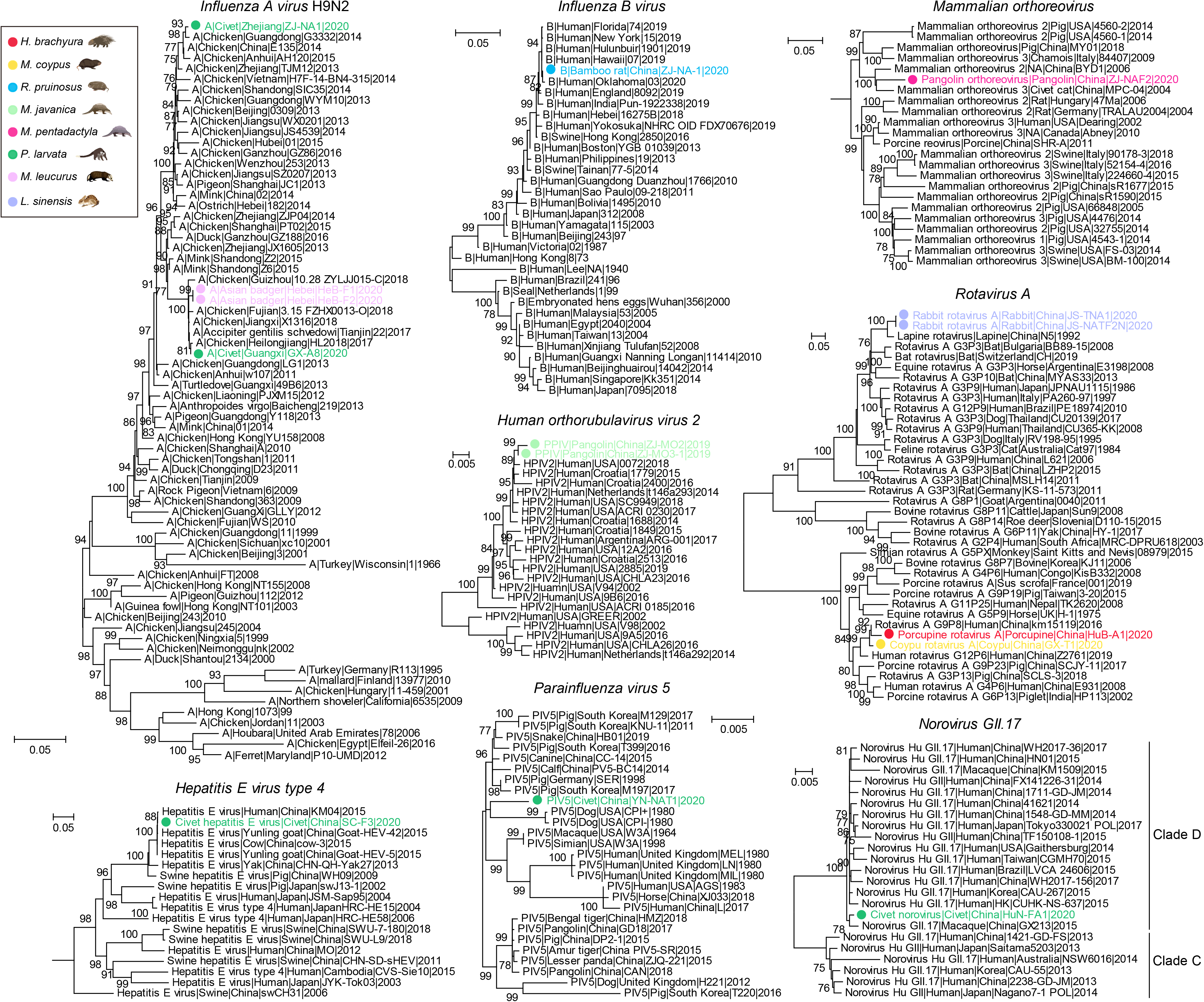
Intra-specific phylogenetic diversity of pathogenic human viruses identified in game animals. Each phylogenetic tree was estimated using a maximum likelihood method based on conserved viral proteins (influenza viruses = HA gene; other RNA viruses = RdRp domain). The trees were midpoint-rooted for clarity only, and the scale bar represents the number of nucleotide substitutions per site. For clarity, only support values > 75% are shown. Within each phylogeny, the names in black represented published viral genomes while the viruses newly identified here are marked by with solid circles and colored by animal hosts as shown in the key. Virus names indicate viral type (subtype), host species, sampling location, strain name and year, from left to right.

We similarly identified cross-species transmission among animal viruses (Figure S1). Bat-associated coronavirus HKU8 was identified in a civet (98.66% amino acid sequence identity at RdRp protein); avian-associated coronavirus HKU17 was identified in porcupines (in both oral and fecal swabs, 94.27% identity); pig associated *Swine pneumovirus 57* was identified in pangolins (98.56% identity), bovine associated *Bovine respirovirus 3* was identified in a rodent species – coypu (98.91%); and multiple *Carnivore protoparvovirus* lineages were identified in both Malayan and Chinese pangolins (96.60-99.89% identity). Interestingly, our data also suggested that *Mangshi virus* (genus *Seadornavirus*, *Reoviridae*), a mosquito-borne virus with unknown vertebrate host but possessing the replication capacity in vertebrate cell lines (i.e. a vector-borne virus), was identified in both porcupine and civet samples. The remaining viruses identified, such as *Rabbit hemorrhagic disease virus* (RHDV), *Aichivirus A* and *Rodent coronavirus* (*Embecovirus*), were more reflective of their known host range.

In addition to existing viral species, we identified 45 previously undescribed putative viral species of vertebrates, the majority of which belonged to the families *Picornaviridae* (n=16), *Astroviridae* (n=9), and *Flavivirdae* (n=9), with others belonging to the *Reoviridae* (*Coypu rotavirus*, *Coypu rotavirus B* and *Civet rotavirus D*), *Tobaniviridae* (*Porcupine nidovirus*), and *Caliciviridae* (*Asian badger vesivirus* and *Bamboo rat sapovirus*) (Figure 3). Interestingly, some of the newly discovered viruses or virus groups, including *Porcupine pegivirus*, *Pangolin pestivirus*, *Bamboo rat sapovirus*, *Mammalian enterovirus*, and civet and coypu *kobuviruses*, had both high prevalence and abundance in their respective host species (Figure 2) and sometimes formed species-specific clusters (Figures 3 and 4). However, their disease manifestation and potential threat to humans is unclear.

### The virome of diseased game animals

We further characterized the virome of sick and/or deceased animals, often using mixed tissue samples. For those animals with a clear record of clinical symptoms, the majority (13/26) were infected with at least one virus species, while others (6/26) were infected with two or more viruses (Figure 5A). Specifically, *Influenza A virus* H9N2 was detected in both Asian badgers presenting with respiratory symptoms; different combinations of *Pangolin hunnivirus*, *Human orthorubulavirus 2*, *Swine pneumovirus 57*, and a diverse group of pangolin pestiviruses were detected in Malayan pangolins suffering from gastroenteritis, pneumonia and multiple organ hemorrhage; *Bat coronavirus* HKU8 and *Norovirus GII.17* were simultaneously detected in a civet with diarrheal symptoms; and a novel *pegivirus* (*Porcupine pegivirus*) was detected in two Malayan porcupines exhibiting depressive-like behavior, paralysis and trichomadesis symptoms. Conversely, no viral pathogens were detected in some deceased animals that had no recorded clinical symptoms and hence an unknown cause of death, including masked civets (n=5), Asian badger (n=1), Chinese ferret-badger (n=1), hog badger (n=1), Pallas’s squirrel (n=1), Chinese pangolin (n=1) and Malayan pangolin (n=1). In addition to diseased animals, pathogens were also detected in seemingly healthy animals (Figure 5B). For example, *Influenza A virus* H9N2 was identified in a healthy civet, although the same virus may result in influenza-like symptoms in Asian badgers. Similarly, *Hepatitis E virus*, *Mangshi virus*, *Parainfluenza virus 5* and *Feline coronavirus* were detected in healthy civets, and a common human pathogen, *Influenza B virus*, was detected in a seemingly healthy bamboo rat population (Figure 5B).

**Figure 5.**
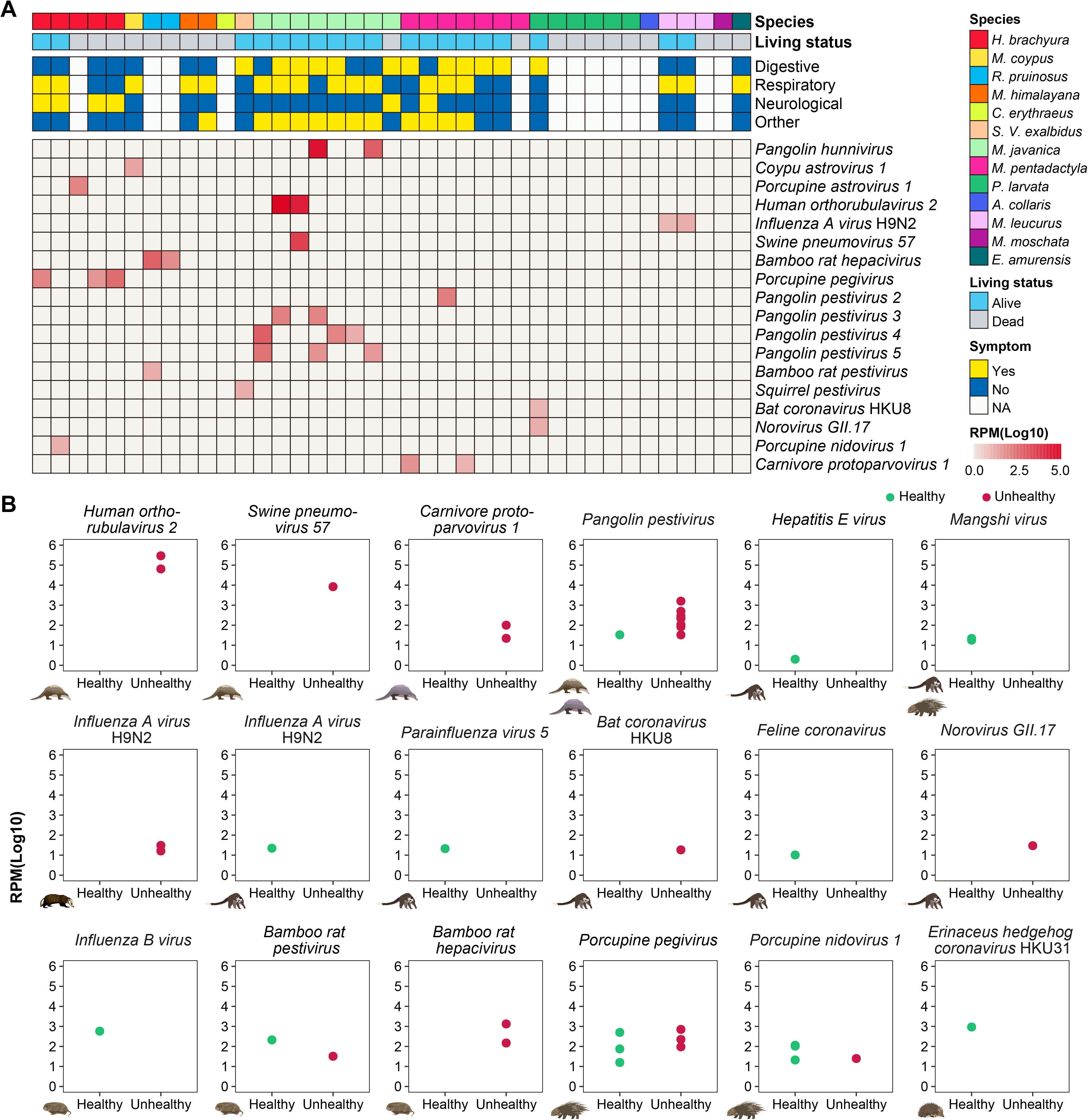
Virome characterization in diseased game animals. (A) The distribution and abundance of vertebrate-associated viruses in diseased game animals. The relative abundance of each virus in each library was calculated and normalized by the number of mapped reads per million total reads (RPM). To reduce the possibility of contamination, only RPM > 10 are shown. Disease symptoms were recorded when sampling and classified into digestive (intestine and stomach related), respiratory (lung related and influenza-like), neurological (paralysis, convulsion and trauma) and other (spleen and liver related, injury, weakness, loss of appetite etc.) symptoms. Living status (i.e. dead or alive) was recorded when sampling and depicted by different colors. (B) Abundance comparisons between healthy and unhealthy game animals (see symbols) for 18 pathogenic viruses.

### Characterization of viruses at high emergence potential

Among the 71 vertebrate-associated viruses identified here, we characterized the epidemiological patterns of 18 that we considered to pose a greater risk for infection of humans or other animal species. This risk assessment was simply based on perceived zoonotic potential (i.e. likely ability to infect humans) and/or tendency to jump species barriers and infect other animal hosts (Figure 6A). Most of these putative high-risk viruses exhibited a strong geographic structure, clustering according to Chinese province. For example, *Bat coronavirus* HKU8 was found in Hunan province, *Bovine respirovirus 3* was found in Hebei province, and *Influenza B virus* was found in Zhejiang province, while *Parainfluenza virus 5* was sampled in Yunnan province (Figure 6B). In contrast, several viruses had a wider geographical range. For example, *Influenza A virus* H9N2 was identified in Hebei province (northern China) and Guangxi province (southern China), while *Rotavirus A* was identified in Hubei province (central China), Jiangxi province (eastern China), and Guangxi province (southern China) (Figure 6B).

**Figure 6.**
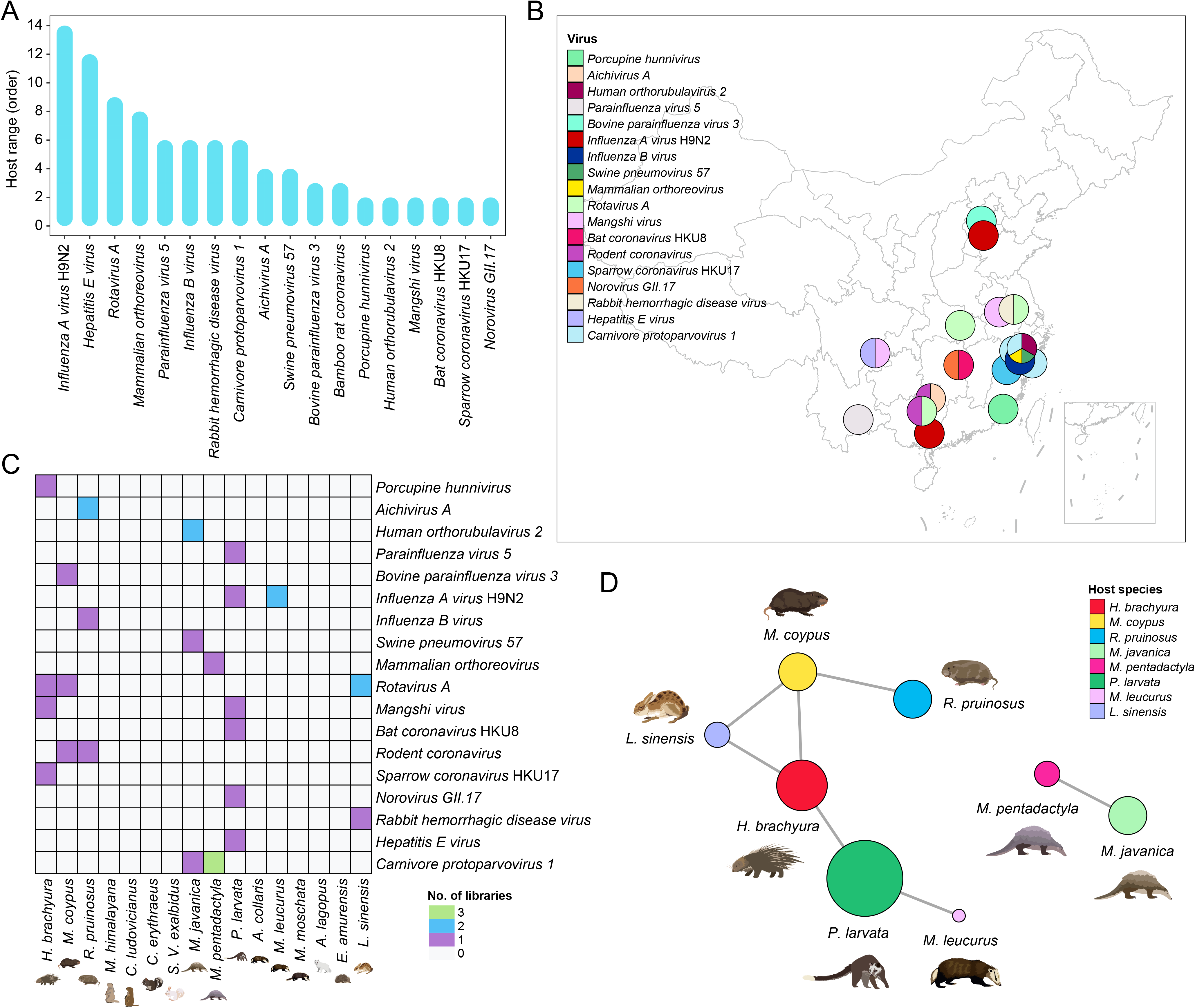
Epidemiological patterns of “high-risk” viruses with the potential to infect humans or other mammals. (A) Host range of potential zoonotic viruses as indicated by the level of the number of mammalian orders infected. For known viruses, a statistical analysis of host range was conducted based on host species information retrieved from NCBI and expanded with the hosts newly identified here. For unknown viruses, a statistical analysis of host range was conducted based only on the host species information obtained in this study. (B) Geographical locations of potential zoonotic viruses identified in this study. Any viruses shared between more than two hosts were included and regarded as potentially zoonotic viruses. (C) Distribution of potential zoonotic viruses in each host species. (D) Associations between potential zoonotic viruses and game species hosts. The size of the colored circles indicates the number of potential zoonotic viral species carried by each host species, while the thickness of the line indicates number of potential zoonotic viral species shared by each host species.

Several viruses exhibited high prevalence as well as signatures of cross-species transmission, appearing in multiple libraries within the same or different species. For example, *Rotavirus A* was detected in three animal species (Figure 6C), while *Mangshi virus* was discovered in both porcupines and civets, although human infection has not yet been identified.

Finally, we compared the likelihood of different game animals carrying pathogens of greater potential zoonotic risk and the presence of these pathogens among different species (Figure 6D). Among the animals studied, civets carried the highest number of potentially high-risk viruses (n=6), followed by porcupines (n=4), coypus (n=3), bamboo rats (n=3) and Malayan pangolins (n=3). Hence, there is a risk of cross-species transmission to other groups of game animals, and it is notable that Malayan and Chinese pangolins were not surveyed for viruses until very recently.

## Discussion

We performed a large-scale survey of game animals that are commonly hunted or consumed for food in China, among which many species were examined for the first time and some have been banned by the Chinese Government for trading or artificial breeding since the onset of the COVID-19 pandemic. This analysis identified a wide diversity of both previously described and novel virus species, some of which may pose direct threat to human health. Notably, we did not identify any SARS-CoV-2-like or SARS-CoV-like sequences. This included in Malayan pangolins from which SARS-CoV-2-like viruses have previously been identified^11,12,24^. One explanation for the lack of pangolin SARS-like viruses in our samples is that previous virus-positive samples were all confiscated by the custom authorities in Guangdong and Guangxi provinces^11,12,24,37^, both of which were characterized by an intensive network of animal smuggling^38,39^. In contrast, our pangolin samples were obtained from Zhejiang province in eastern China which does not have an international border that would facilitate smuggling. Clearly, the ongoing surveillance of pangolins for zoonotic pathogens is of utmost importance.

Although no SARS-like viruses were identified in this study, we did identify a clear cross-species transmission event involving a bat-associated coronavirus that might seed a disease outbreak. Specifically, we identified a bat-associated virus - BatCoV-HKU8 – in a civet that represents a different mammalian order (Carnivora). Genomic comparisons revealed a high sequence identity across most of the virus genome (>93.84% amino acid identity) with the exception of the N-terminal proportion of the S1 subunit of the spike protein (36.52-81.74% amino acid identity) that contained a receptor-binding domain (RBD) that might be associated with host adaptation. In addition, bird-to-porcupine (i.e., *Sparrow coronavirus HKU17*) and bat-to-hedgehog (*Erinaceus amurensis hedgehog coronavirus HKU31*, Ea-HedCoV HKU31) spill-overs or host switches were also identified within the *Coronaviridae*. The former was also related to a swine virus (Porcine deltacoronavirus) recently identified in three Haitian children with acute undifferentiated febrile illness^40^. The latter clustered within the MERS-like virus group (Figure 3) that have previously been identified in Amur hedgehogs and European hedgehogs^41,42^, suggesting that hedgehogs are likely important reservoirs for MERS-like viruses. Collectively, we suggest that the cross-species transmission from bats or avian hosts to game animals may represent a possible pathway for the virus to move from bats to humans. Indeed, bat-associated alpha- and beta-coronaviruses are notorious for being the ancestors for those viruses that cause disease outbreaks in humans, including SARS-CoV^43,44^, SARS-CoV-2^34^, MERS-CoV^45,46^, HCoV-229E^47,48^ and HCoV-NL63^49^, although they commonly pass through so-called “intermediate” hosts such as civets and raccoon dogs for SARS-CoVs^10^, camels for MERS-CoVs^50^, and alpacas for HCoV-229E^51^. As such, our study underlines the fact that coronaviruses are subject to relatively frequent host shifts from reservoir hosts to terrestrial mammals and therefore pose a direct threat to wildlife animal handlers or during food consumption.

Another virus that might pose an immediate threat to human health is avian influenza virus H9N2 that has increased in prevalence in live poultry markets workers in China^52^. H9N2 has gradually replaced H5N6 and H7N9 as the most prevalent AIV subtype in both chickens and ducks^53^ and has caused numerous human infections in China. Indeed, in comparisons to other currently circulating avian influenza viruses, H9N2 has clear potential to infect humans because almost all subtype H9 AIVs possess human-type receptor-binding ability^54,55^. Interestingly, the H9N2 virus identified here exhibited the 155T, 183N and 226L amino substitutions in the HA1 receptor-binding domain (RBD) previously associated with human infection^56^. It is therefore of considerable significance that we detected H9N2 in two game animal species - Asian badgers and a civet from northern and southern China, respectively (Figure 6A and 6B). Furthermore, Asian badgers displayed obvious respiratory symptoms, suggesting a potential respiratory transmission route that increases the risk of disease transmission to humans.

Our study also revealed that game animals are important hosts for viruses that are related to diseases in human and/or domestic animals. Not only did we sample common zoonotic pathogens such as *Rotavirus A*, *hepatitis E virus*, and *Mammalian orthoreovirus, Parainfluenza virus 5* that are known be harbored by a wide range of mammalian hosts, but also viruses previously thought to be specific to certain mammalian host groups. For example, we present the first evidence of human pathogens such as *Influenza B virus*, *Human parainfluenza virus 2*, and *Norovirus GII.17* in bamboo rats, Malayan pangolins, and civets, indicative of human-to-wildlife transmission. Of note, *Influenza B viruses* have previously been identified in other mammalian species including seals and pigs^57,58^. Similarly, BatCoV-HKU8, *Bovine respirovirus 3*, *Carnivore protoparvovirus* and *Influenza A virus* H9N2, which are predominantly found in bats, cattle, carnivores, and avian hosts, respectively, were now detected in a wide range of game animals from different mammalian orders to their perceived reservoir host groups. It is currently unclear how these viruses are maintained in these game animal species compared to other hosts. Nevertheless, their capability to carry these viruses, even for a short period of time, may contribute to the virus transmission chains or the emergence of new variants. Indeed, the potential for game animals to act as a separate transmission chain might in part explain the sometimes sudden and unexpected emergence of new virus variants in humans or domestic animals, as observed in several norovirus outbreaks^59^. In addition, our results show that these viruses were present in seemingly healthy animals and that there is ongoing transmission among different species of game animals (Figure 5). This will increase the likelihood of these animals acting as intermediate hosts and again highlights the risk of close contact with game animals.

More broadly, our results provide important insights to those game animals and their viruses that might lead to the next pandemic or epizootic. Indeed, the viruses discovered in this study can be divided into two categories – those with restricted host range and those with the capability to infect animals from different mammalian orders and hence that seem able to overcome host genetic barriers. The latter category merits most attention as they underpin the great majority of human epidemics and pandemics.

## Material and methods

### Sample collection

Sampling of game animals was performed between June 2017 and June 2021 at locations representing the natural habitat of animals, artificial breeding sites and zoos across 19 provinces in China. Accordingly, a total of 2595 samples from 1726 animal individuals and 16 species were obtained. These comprised: *Hystrix brachyura* (Malayan porcupine, n=379), *Myocastor coypus* (coypu, n=300), *Rhizomys pruinosus* (bamboo rat, n=349), *Marmota himalayana* (himalayan marmot, n=19), *Cynomys ludovicianus* (prairie dog, n=60), *Callosciurus erythraeus* (Pallas’s squirrel, n=1), *Sciurus vulgaris exalbidus* (Siberian red squirrel, n=1) from the order *Rodentia*; *Manis javanica* (Malayan pangolin, n=20), *Manis pentadactyla* (Chinese pangolin, n=12) from the order *Pholidota*; *Arctonyx collaris* (hog badger, n=1), *Meles leucurus* (Asian badger, n=54), *Melogale moschata* (Chinese ferret-badger, n=1), *Paguma larvata* (civet, n=423),*Alopex lagopus* (Arctic fox, n=1) from the order *Carnivora*; *Erinaceus amurensis* (Amur hedgehog, n=24) from the order *Eulipotyphla*; and *Lepus sinensis* (Chinese hare, n=71) from the order *Lagomorpha* (Figure 1A and 1B). Pharyngeal, anal, nasal and fecal swabs were obtained from living animals, whereas tissue samples (heart, liver, spleen, lung, kidney, intestine, stomach) were obtained from deceased animals. All animal specimens were kept in dry ice before transfer to a −80° freezer for storage.

Host species information was initially identified by experienced field biologists upon capture based on morphological characteristics, then further confirmed by Sanger sequencing of the cytochrome b (*CytB*) gene^60^. The procedures for sampling and sample processing were approved by the ethics committee of Institute of Military Veterinary Medicine, Academy of Military Medical Sciences, Academy of Military Sciences (No. IACUC of AMMS-11-2020-012).

### RNA extraction, library preparation and sequencing

Samples were pooled according to the species, location, health condition, feeding (living) condition, for subsequent RNA extraction and library construction. The total RNA from each pool was extracted using TRIzol® Reagent according to the manufacturer’s instructions (Invitrogen). Genomic DNA was subsequently removed using DNase I (Takara). Meta-transcriptome library preparation of each pool was carried out using the TruSeq™ Stranded Total RNA Sample Preparation Kit from Illumina (San Diego, CA) after removal of host ribosomal RNA using Illumina Ribo-Zero™ rRNA Removal Kits (San Diego, CA). Paired-end (150 bp) sequencing of each RNA library was performed using the Illumina Novaseq 6000 platform.

### Virus discovery and confirmation

For each library, sequencing reads were first quality controlled using bbduk.sh (https://sourceforge.net/projects/bbmap/; parameters, maq=10 qtrim=r trimq=10 ftl=1 minlen=90). The remaining reads were assembled *de novo* using MEGAHIT^61^ (version 1.2.6) deploying default parameters. The assembled contigs were compared against the NCBI non-redundant protein database (nr) using Diamond blastx (version 0.9.25.26)^62^. The *E*-value cut-off was set at 1*E*-3 to maintain high sensitivity at a low false-positive rate. Taxonomic lineage information was obtained for the top blast hit of each contig, and those classified under kingdom “Viruses” were identified as probable virus hits. Contigs with unassembled overlaps were merged using the SeqMan program implemented in the Lasergene software package 7.1 (DNAstar)^63^. The final virus genomes were verified by mapping reads to the corresponding contigs and inspecting the mapping results using Geneious^64^. The presence of viruses was further verified using RT-PCR and nested RT-PCR on the original RNA samples using primer sets designed based on the viral genome sequences. The PCR products were then sequenced and compared with the original template. For incomplete viral genomes, the internal gaps were filled by RT-PCR and Sanger sequencing, whereas the genome termini were determined using RACE analyses.

Among the virus contigs described here, those likely associated with vertebrates, specifically vertebrate-specific viruses and vector-borne vertebrate viruses, were preliminarily identified based on the taxonomic lineage information of the blastx results and confirmed by phylogenetic analyses. Those grouping within a vertebrate-specific or vector-borne virus family/genus/group were subsequently identified as vertebrate viruses. We employed a simplified criterion of <80% nucleotide identity at the whole viral genome level to assign new virus species which were then verified by detailed phylogenetic analysis (see below).

### Estimation of virus prevalence and abundance

Each virus sequence was first assigned to specific virus species based on the level of sequence similarity to each other and to reference sequences. To estimate the relative abundance of each virus species in each library, quality-trimmed reads were firstly mapped to the SILIVA database (www.arb-silva.de, version 132.1) using Bowtie2 (version 2.3.5.1)^65^ to remove reads associated with ribosomal RNA. The unmapped reads were subsequently mapped to confirmed viral genomes using the “end-to-end” setting, and the abundance of each virus species was estimated as the number of mapped reads per million total reads (RPM) in each library. To limit false-positives, we set the RPM greater than or equal to 10 as providing evidence for a positive virus hit.

### Evolutionary analysis

To infer the evolutionary history of all vertebrate-associated viruses here, we first assigned these viral sequences into their respective (vertebrate associated) viral family or genus based on the Diamond blastx results. The amino acids of viral replicase proteins (i.e. RNA viruses = RdRp domain; DNA viruses = major capsid protein) of these virus genomes were then aligned with those of related viruses downloaded from GenBank using MAFFT^66^ (version 7.475), employing the L-INS-I algorithm. Following sequence alignment, all ambiguous aligned regions were removed using TrimAL^67^ (version 1.2). Phylogenetic trees were then estimated for the sequence alignment of each family using the maximum likelihood method available in PhyML (version 3.1)^68^, employing the LG model of amino acid substitution and a subtree pruning and the regrafting (SPR) branch swapping algorithm. A similar procedure was utilized for intra-specific virus phylogenetic trees, with the exception that the greater sequence identity meant that genome-scale nucleotide sequences were used in sequence alignments and a general-time reversible (GTR) substitution model employed for phylogenetic inference.

---

**Figure S1. Intra-specific phylogenetic diversity of non-human pathogenic viruses identified in game animals.** Each phylogenetic tree was estimated using a maximum likelihood method based on conserved viral proteins (RNA viruses = RdRp domain; DNA viruses = Major capsid protein). The trees were midpoint-rooted with the scale bar denoting the number of nucleotide substitutions per site. For clarity, only bootstrap values > 75% are shown. Within each phylogeny, virus names in black represent published viral genomes while those newly identified here are marked by with solid circles and colored by animal host (see animal symbols in the legend bar). Virus names indicate viral type (subtype), host species, sampling location, strain name and year, from left to right. For HKU8 and the rodent coronavirus we specified the host as detailed as by the common name or Latin name.

## References

1. Huong, N. Q. et al. Coronavirus testing indicates transmission risk increases along wildlife supply chains for human consumption in Viet Nam, 2013-2014. PLoS One 15, e0237129, doi:10.1371/journal.pone.0237129 (2020).

2. Philavong, C. et al. Perception of health risks in Lao market vendors. Zoonoses Public Health 67, 796–804, doi:10.1111/zph.12759 (2020).

3. Shivaprakash, K. N., Sen, S., Paul, S., Kiesecker, J. M. & Bawa, K. S. Mammals, wildlife trade, and the next global pandemic. Curr Biol 31, 3671–3677 e3673, doi:10.1016/j.cub.2021.06.006 (2021).

4. Damania, B. R. An Economic Assessment of Wildlife Farming and Conservation. Conservation Biology 19, 1222–1233 (2005).

5. Xiao, X., Newman, C., Buesching, C. D., Macdonald, D. W. & Zhou, Z. M. Animal sales from Wuhan wet markets immediately prior to the COVID-19 pandemic. Sci Rep 11, 11898, doi:10.1038/s41598-021-91470-2 (2021).

6. Holmes, E. C. et al. The origins of SARS-CoV-2: A critical review. Cell 184, 4848–4856, doi:10.1016/j.cell.2021.08.017 (2021).

7. Aditya, V., Goswami, R., Mendis, A. & Roopa, R. Scale of the issue: Mapping the impact of the COVID-19 lockdown on pangolin trade across India. Biol Conserv 257, 109136, doi:10.1016/j.biocon.2021.109136 (2021).

8. Sun, J. et al. COVID-19: Epidemiology, Evolution, and Cross-Disciplinary Perspectives. Trends Mol Med 26, 483–495, doi:10.1016/j.molmed.2020.02.008 (2020).

9. Skowronski, D. M. et al. Severe acute respiratory syndrome (SARS): a year in review. Annu Rev Med 56, 357–381, doi:10.1146/annurev.med.56.091103.134135 (2005).

10. Guan, Y. et al. Isolation and characterization of viruses related to the SARS coronavirus from animals in southern China. Science 302, 276–278, doi:10.1126/science.1087139 (2003).

11. Lam, T. T. et al. Identifying SARS-CoV-2-related coronaviruses in Malayan pangolins. Nature 583, 282–285, doi:10.1038/s41586-020-2169-0 (2020).

12. Xiao, K. et al. Isolation of SARS-CoV-2-related coronavirus from Malayan pangolins. Nature 583, 286–289, doi:10.1038/s41586-020-2313-x (2020).

13. Zhang, T., Wu, Q. & Zhang, Z. Probable Pangolin Origin of SARS-CoV-2 Associated with the COVID-19 Outbreak. Current Biology 30, 1346–1351.e1342, doi:10.1016/j.cub.2020.03.022 (2020).

14. Chen, J. P., Andersen, D. H., Veron, G., Randi, E. & Zhang, S. Y. Isolation and characterization of polymorphic microsatellite markers for the masked palm civet (Paguma larvata). Biochem Genet 46, 392–397, doi:10.1007/s10528-008-9157-7 (2008).

15. Martine Peeters, V. C., Bernadette Abela, Philippe Auzel, Xavier Pourrut, Frederic Bibollet-Ruche, Severin Loul, Florian Liegeois, Cristelle Butel, Denis Koulagna, Eitel Mpoudi-Ngole, George M Shaw, Beatrice H Hahn, Eric Delaporte. Risk to human health from a plethora of simian immunodeficiency viruses in primate bushmeat. Emerg Infect Dis 8, 451–457, doi:10.3201/eid0805.010522 (2002).

16. Locatelli, S. & Peeters, M. Cross-species transmission of simian retroviruses: how and why they could lead to the emergence of new diseases in the human population. AIDS 26, 659–673, doi:10.1097/QAD.0b013e328350fb68 (2012).

17. Aghokeng, A. F. et al. Extensive survey on the prevalence and genetic diversity of SIVs in primate bushmeat provides insights into risks for potential new cross-species transmissions. Infect Genet Evol 10, 386–396, doi:10.1016/j.meegid.2009.04.014 (2010).

18. Techangamsuwan, S. et al. Pathologic and Molecular Virologic Characterization of a Canine Distemper Outbreak in Farmed Civets. Vet Pathol 52, 724–731, doi:10.1177/0300985814551580 (2015).

19. Wang, X. et al. Complete Genome Sequence of Parainfluenza Virus 5 (PIV5) from a Sunda Pangolin (Manis javanica) in China. J Wildl Dis 55, 947–950 (2019).

20. Wang, S. L. et al. Fatal canine parvovirus-2 (CPV-2) infection in a rescued free-ranging Taiwanese pangolin (Manis pentadactyla pentadactyla). Transbound Emerg Dis 67, 1074–1081, doi:10.1111/tbed.13469 (2020).

21. Dai, X., Shang, G., Lu, S., Yang, J. & Xu, J. A new subtype of eastern tick-borne encephalitis virus discovered in Qinghai-Tibet Plateau, China. Emerg Microbes Infect 7, 74, doi:10.1038/s41426-018-0081-6 (2018).

22. Kent, A. et al. Genital Tract Screening Finds Widespread Infection with Mustelid Gammaherpesvirus 1 in the European Badger ( Meles Meles). J Wildl Dis 54, 133–137, doi:10.7589/2016-12-274 (2018).

23. Banks, M., King, D. P., Daniells, C., Stagg, D. A. & Gavier-Widen, D. Partial characterization of a novel gammaherpesvirus isolated from a European badger (Meles meles). J Gen Virol 83, 1325–1330, doi:10.1099/0022-1317-83-6-1325 (2002).

24. Liu, P., Chen, W. & Chen, J. P. Viral Metagenomics Revealed Sendai Virus and Coronavirus Infection of Malayan Pangolins (Manis javanica). Viruses 11, doi:10.3390/v11110979 (2019).

25. Luo, X. L. et al. Marmota himalayana in the Qinghai-Tibetan plateau as a special host for bi-segmented and unsegmented picobirnaviruses. Emerg Microbes Infect 7, 20, doi:10.1038/s41426-018-0020-6 (2018).

26. Ao, Y. et al. Two novel bocaparvovirus species identified in wild Himalayan marmots. Sci China Life Sci 60, 1348–1356, doi:10.1007/s11427-017-9231-4 (2017).

27. Rogier Bodewes, A. R.-G., Claudia M E Schapendonk, Judith M A van den Brand, Albert D M E Osterhaus, Saskia L Smits. Viral metagenomic analysis of feces of wild small carnivores. Virol J 11, doi:10.1186/1743-422X-11-89 (2014).

28. Shao, Y.-H., Han, Z.-X., Chen, L.-F., Kong, X.-G. & Liu, S.-W. J. B. d. x. b. C. J. o. V. Isolation and identification of A reovirus from masked civet cats (Paguma Larvata). Bing Du Xue Bao 24, 376–382 (2008).

29. Abe, M. et al. Molecular characterization of rotaviruses in a Japanese raccoon dog (Nyctereutes procyonoides) and a masked palm civet (Paguma larvata) in Japan. Vet Microbiol 146, 253–259, doi:10.1016/j.vetmic.2010.05.019 (2010).

30. Arnaboldi, S. et al. Hepatitis E Virus (HEV) Spread and Genetic Diversity in Game Animals in Northern Italy. Food Environ Virol 13, 146–153, doi:10.1007/s12560-021-09467-z (2021).

31. Doro, R., Farkas, S. L., Martella, V. & Banyai, K. Zoonotic transmission of rotavirus: surveillance and control. Expert Rev Anti Infect Ther 13, 1337–1350, doi:10.1586/14787210.2015.1089171 (2015).

32. Nimgaonkar, I., Ding, Q., Schwartz, R. E. & Ploss, A. Hepatitis E virus: advances and challenges. Nat Rev Gastroenterol Hepatol 15, 96–110, doi:10.1038/nrgastro.2017.150 (2018).

33. Pavio, N., Meng, X. J. & Doceul, V. Zoonotic origin of hepatitis E. Curr Opin Virol 10, 34–41, doi:10.1016/j.coviro.2014.12.006 (2015).

34. Zhou, P. et al. A pneumonia outbreak associated with a new coronavirus of probable bat origin. Nature 579, 270–273, doi:10.1038/s41586-020-2012-7 (2020).

35. Zhou, H. et al. Identification of novel bat coronaviruses sheds light on the evolutionary origins of SARS-CoV-2 and related viruses. Cell, doi:10.1016/j.cell.2021.06.008 (2021).

36. Wu, D. et al. Civets are equally susceptible to experimental infection by two different severe acute respiratory syndrome coronavirus isolates. J Virol 79, 2620–2625, doi:10.1128/JVI.79.4.2620-2625.2005 (2005).

37. Yang, S. et al. Digging metagenomic data of pangolins revealed SARS-CoV-2 related viruses and other significant viruses. J Med Virol 93, 1786–1791, doi:10.1002/jmv.26524 (2021).

38. Zhang, L., Hua, N. & Sun, S. Wildlife trade, consumption and conservation awareness in southwest China. Biodivers Conserv 17, 1493–1516, doi:10.1007/s10531-008-9358-8 (2008).

39. Huang, X. Q. et al. Prosecution records reveal pangolin trading networks in China, 2014-2019. Zool Res 42, 666–670, doi:10.24272/j.issn.2095-8137.2021.156 (2021).

40. Lednicky, J. A. et al. Emergence of porcine delta-coronavirus pathogenic infections among children in Haiti through independent zoonoses and convergent evolution. medRxiv, doi:10.1101/2021.03.19.21253391 (2021).

41. Corman, V. M. et al. Characterization of a novel betacoronavirus related to middle East respiratory syndrome coronavirus in European hedgehogs. J Virol 88, 717–724, doi:10.1128/JVI.01600-13 (2014).

42. Lau, S. K. P. et al. Identification of a Novel Betacoronavirus (Merbecovirus) in Amur Hedgehogs from China. Viruses 11, doi:10.3390/v11110980 (2019).

43. Thomas G Ksiazek1, D. E., Cynthia S Goldsmith, Sherif R Zaki, Teresa Peret, Shannon Emery, Suxiang Tong, Carlo Urbani, James A Comer, Wilina Lim, Pierre E Rollin, Scott F Dowell, Ai-Ee Ling, Charles D Humphrey, Wun-Ju Shieh, Jeannette Guarner, Christopher D Paddock, Paul Rota, Barry Fields, Joseph DeRisi, Jyh-Yuan Yang, Nancy Cox, James M Hughes, James W LeDuc, William J Bellini, Larry J Anderson, SARS Working Group. A novel coronavirus associated with severe acute respiratory syndrome. N Engl J Med 348, 1953–1966, doi:10.1056/NEJMoa030781 (2003).

44. Rota, P. A. et al. Characterization of a novel coronavirus associated with severe acute respiratory syndrome. Science 300, 1394–1399, doi:10.1126/science.1085952 (2003).

45. Assiri, A. et al. Hospital outbreak of Middle East respiratory syndrome coronavirus. N Engl J Med 369, 407–416, doi:10.1056/NEJMoa1306742 (2013).

46. Zaki, A. M., van Boheemen, S., Bestebroer, T. M., Osterhaus, A. D. & Fouchier, R. A. Isolation of a novel coronavirus from a man with pneumonia in Saudi Arabia. N Engl J Med 367, 1814–1820, doi:10.1056/NEJMoa1211721 (2012).

47. Hamre D Fau - Procknow, J. J. & Procknow, J. J. A new virus isolated from the human respiratory tract. (1996).

48. Ntumvi, N. F. et al. Wildlife in Cameroon harbor diverse coronaviruses including many isolates closely related to human coronavirus 229E. bioRxiv, doi:10.1101/2021.09.03.458874 (2021).

49. van der Hoek, L., Pyrc, K. & Berkhout, B. Human coronavirus NL63, a new respiratory virus. FEMS Microbiol Rev 30, 760–773, doi:10.1111/j.1574-6976.2006.00032.x (2006).

50. Dudas, G., Carvalho, L. M., Rambaut, A. & Bedford, T. MERS-CoV spillover at the camel-human interface. Elife 7, doi:10.7554/eLife.31257 (2018).

51. Crossley, B. M., Mock, R. E., Callison, S. A. & Hietala, S. K. Identification and characterization of a novel alpaca respiratory coronavirus most closely related to the human coronavirus 229E. Viruses 4, 3689–3700, doi:10.3390/v4123689 (2012).

52. Li, X. et al. A comprehensive retrospective study of the seroprevalence of H9N2 avian influenza viruses in occupationally exposed populations in China. PLoS One 12, e0178328, doi:10.1371/journal.pone.0178328 (2017).

53. Bi, Y. et al. Dominant subtype switch in avian influenza viruses during 2016-2019 in China. Nat Commun 11, 5909, doi:10.1038/s41467-020-19671-3 (2020).

54. Bi, Y. et al. Genesis, Evolution and Prevalence of H5N6 Avian Influenza Viruses in China. Cell Host Microbe 20, 810–821, doi:10.1016/j.chom.2016.10.022 (2016).

55. Liu, D., Shi, W. & Gao, G. F. Poultry carrying H9N2 act as incubators for novel human avian influenza viruses. The Lancet 383, doi:10.1016/s0140-6736(14)60386-x (2014).

56. Li, X. et al. Genetics, receptor binding property, and transmissibility in mammals of naturally isolated H9N2 Avian Influenza viruses. PLoS Pathog 10, e1004508, doi:10.1371/journal.ppat.1004508 (2014).

57. Osterhaus, A. D., Rimmelzwaan, G. F., Martina, B. E., Bestebroer, T. M. & Fouchier, R. A. Influenza B virus in seals. Science 288, 1051–1053, doi:10.1126/science.288.5468.1051 (2000).

58. Ran, Z. et al. Domestic pigs are susceptible to infection with influenza B viruses. J Virol 89, 4818–4826, doi:10.1128/JVI.00059-15 (2015).

59. Villabruna, N., Koopmans, M. P. G. & de Graaf, M. Animals as Reservoir for Human Norovirus. Viruses 11, doi:10.3390/v11050478 (2019).

60. Guillén-Servent, A. & Francis, C. M. A new species of bat of the Hipposideros bicolor group (Chiroptera: Hipposideridae) from Central Laos, with evidence of convergent evolution with Sundaic taxa. Acta Chiropterologica 8, 39–61, doi:10.3161/1733-5329(2006)8 [39:Ansobo]2.0.Co;2 (2006).

61. Li, D., Liu, C.-M., Luo, R., Sadakane, K. & Lam, T.-W. MEGAHIT: an ultra-fast single-node solution for large and complex metagenomics assembly via succinct de Bruijn graph. Bioinformatics (Oxford, England) 31, 1674–1676, doi:10.1093/bioinformatics/btv033 (2015).

62. Buchfink, B. A.-O. X., Reuter, K. A.-O. & Drost, H. A.-O. X. Sensitive protein alignments at tree-of-life scale using DIAMOND. Nat Methods (2021).

63. Clewley, J. P. Macintosh sequence analysis software. DNAStar’s LaserGene. Molecular biotechnology 3, 221–224 (1995).

64. Kearse, M. et al. Geneious Basic: an integrated and extendable desktop software platform for the organization and analysis of sequence data. Bioinformatics 28, 1647–1649, doi:10.1093/bioinformatics/bts199 (2012).

65. Langmead, B. & Salzberg, S. L. Fast gapped-read alignment with Bowtie 2. Nature methods 9, 357–359, doi:10.1038/nmeth.1923 (2012).

66. Katoh, K. & Standley, D. M. MAFFT multiple sequence alignment software version 7: improvements in performance and usability. Molecular biology and evolution 30, 772–780, doi:10.1093/molbev/mst010 (2013).

67. Capella-Gutiérrez, S., Silla-Martínez, J. M. & Gabaldón, T. trimAl: a tool for automated alignment trimming in large-scale phylogenetic analyses. Bioinformatics (Oxford, England) 25, 1972–1973, doi:10.1093/bioinformatics/btp348 (2009).

68. Guindon, S. & Gascuel, O. A simple, fast, and accurate algorithm to estimate large phylogenies by maximum likelihood. Systematic biology 52, 696–704 (2003).

