## Supplementary figures and images for "Total virome characterizations of game animals in China reveals a spectrum of emerging viral pathogens"

### Supplemental Figure 1

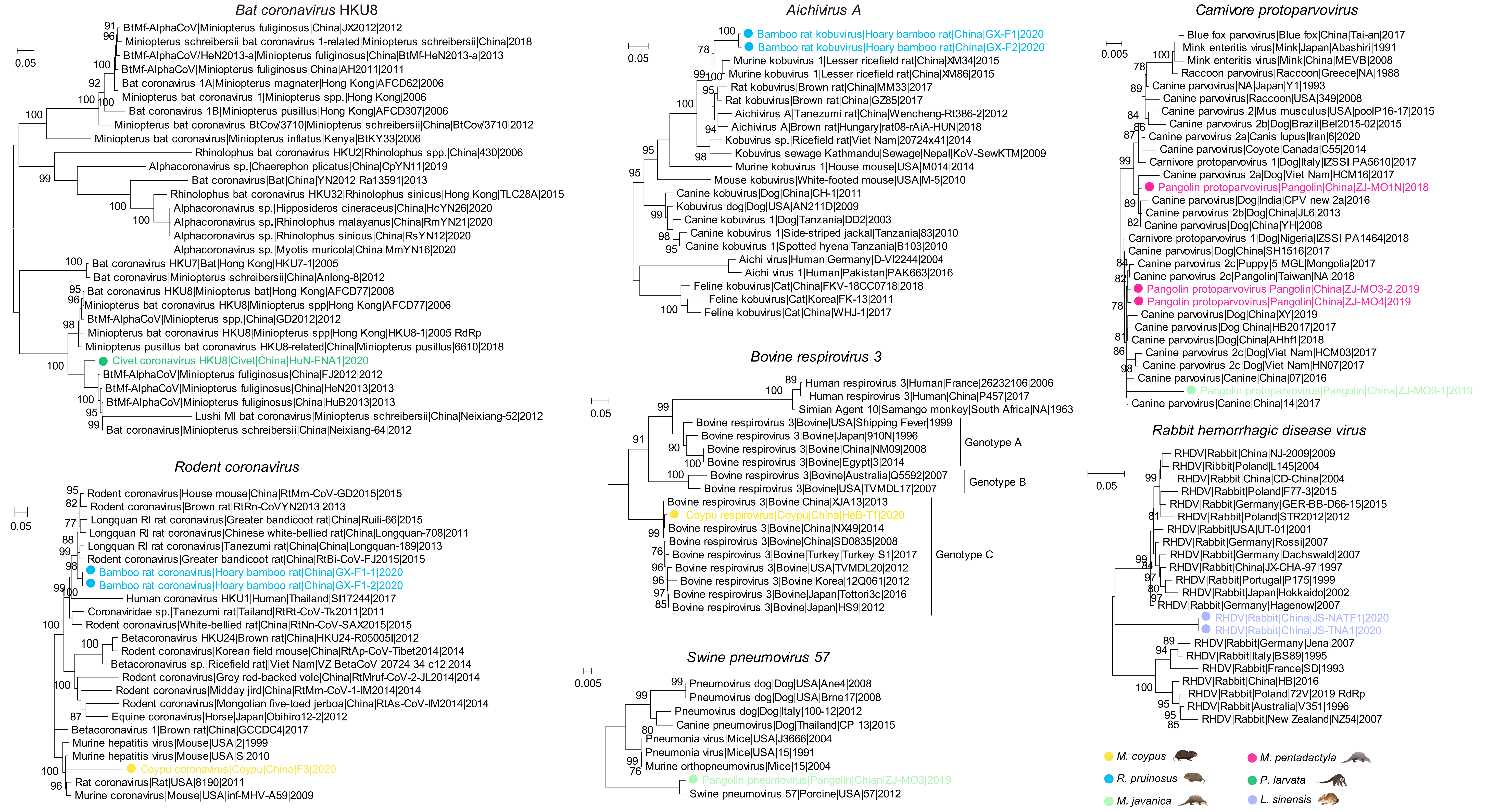
